# Impact of temperature on vector competence of *Culex pipiens molestus*: implications for Usutu virus transmission in temperate regions

**DOI:** 10.1101/2025.04.29.651192

**Authors:** Nicola Seechurn, Jack Pilgrim, Ken Sherlock, Jolanta Tanianis-Hughes, Marcus Blagrove, Grant L. Hughes, Nicholas Johnson, Matthew Baylis

## Abstract

**Introduction:** Usutu virus (USUV) has been detected annually in the southeast of England since 2020. USUV RNA has been identified in wild birds and mosquito populations, and exposure of captive birds to USUV at Zoological Society of London (ZSL) has also been confirmed. Since its first detection in London, USUV’s distribution has expanded across the South East, highlighting a need to understand the transmission dynamics of this virus in the UK. The primary vectors of USUV in the UK are likely *Culex pipiens* mosquitoes. Two biotypes have been identified, the bird-feeding *Cx. pipiens pipiens* and *Cx. pipiens molestus* which shows no restriction in host preference. The latter may play an important role in transmitting USUV from birds to humans.

**Methods:** A laboratory colony of *Cx. pipiens molestus* mosquitoes were orally infected with the London strain of USUV and, incubated at 22 °C, 20 °C and 18° C for up to 28 days. Body samples and mosquito saliva samples were collected and analysed using a quantitative real-time reverse transcription PCR to determine infection and transmission potential, respectively.

**Results:** USUV RNA was detected in all sample times at all temperatures assessed; with the highest temperature (22 °C) showing the greatest proportion of saliva and body positive samples. At this temperature, there was also an eight-fold increase in the relative viral copy number in the mosquito bodies, which was unobserved at other experimental temperatures. When a more sensitive PCR assay was used at the lowest experimental temperature used (18 °C) USUV RNA was present in the mosquito saliva and body samples for longer and showed a greater proportion of positive samples when compared to 20 °C.

**Conclusion:** This study has demonstrated that *Cx. pipiens molestus* may be able to transmit USUV at 22 °C. Active replication of USUV was identified in the mosquito bodies at 22 °C but could not be demonstrated at lower temperatures, suggesting that 20 °C to 22 °C may be an important temperature threshold in USUV replication and transmission. Utilisation of a more sensitive assay for the lower experimental temperatures revealed that USUV was detectable at 18 °C. Therefore, when conducting infection studies on temperate mosquito-borne viruses, it is important to consider assay sensitivity.

## Background

In the summer of 2020, USUV was first identified in five Eurasian blackbirds (*Turdus merula*) and one house sparrow (*Passer domesticus*) in the southeast of England. Identification of USUV in UK non-migratory bird species, such as house sparrows and Eurasian blackbirds suggests autochthonous transmission by UK endemic mosquitoes [1]. Further surveillance in this area in 2021 identified the presence of USUV in another Eurasian blackbird, suggesting overwintering of USUV and possible further autochthonous transmission [2]. USUV was later detected in rural Cambridgeshire in 2023, demonstrating expansion of the USUV distribution since its first identification [3]. USUV has become endemic in northern and central Europe, however in these countries, summer daily temperatures tend to be higher than in the UK. Repeated detection of USUV over multiple years suggests that UK summer temperatures are permissive. The aim of this work is to determine a threshold at which replication and potential transmission can occur by UK native mosquitoes.

*Culex pipiens* s.l. is the main European USUV vector and is a species complex comprised of *Culex pipiens pipiens, Culex pipiens molestus* and hybrid forms. *Culex pipiens pipiens* is widely distributed and highly abundant across the UK and is the predominant enzootic vector of USUV through circulation of virus between avian and *Culex* mosquito populations [4]. By contrast, *Cx. pipiens molestus* is a voracious human-biting biotype with a distribution mostly limited to underground locations, including the London Underground [5]. This biotype is a potential bridge vector of USUV, being able to spread the virus from birds to humans. Where *Cx. pipiens pipiens* and *Cx. pipiens molestus* have occurred sympatrically in surface habitats, hybridisation can occur [6, 7]. These hybrid species have been described as opportunistic feeders and do not have strict host preferences, leading to their potential as additional bridge vectors [8].

Vector competency for USUV has been evidenced in multiple mosquito species. *Cx. torrentium*; *Cx. quinquefasciatus*; *Cx. restuans*; *Cx. neavei*; *Cx. pipiens molestus*; *Cx. pipiens* s.l.; *Cx. pipiens pipiens*; *Aedes japonicus japonicus*; and *Aedes albopictus* [9-13]. A previous infection study using a UK-derived laboratory colony of USUV demonstrated infection, dissemination and transmission in a single individual mosquito infected with the African reference strain of USUV, suggesting a strong infection barrier to UK Culex mosquitoes for this strain. Successful dissemination and transmission has been demonstrated in other vector competency studies under different experimental conditions [10-12]. Additionally, levels of vector competency have been shown to vary between the two *Culex pipiens* s.l. biotypes, *Cx pipiens pipiens* and *Culex pipiens molestus* [14].

Given the gap in knowledge of vector competence of USUV in cooler regions, we aimed to investigate the effect of vector competency of *Cx. pipiens molestus* under temperate conditions. According to climate data collected by the Met office, average July temperatures have reached 20 °C historically, therefore temperatures above and below this figure were used as the incubation temperatures [15]. The first study (‘Study 1’) was conducted at 22 °C and 20 °C and aimed to assess the vector competency of a laboratory colony of *Cx. pipiens molestus* at these temperatures. The second study (‘Study 2’) aimed to determine if virus was detectable below 20 °C when assay sensitivity is increased. Overall, this infection study assesses the vector competency and importance of assay sensitivity in a laboratory colony of *Cx. pipiens molestus*, incubated between 18 °C and 22 °C, using the USUV London strain of the African 3 lineage [1].

## Methods

### Mosquito rearing and virus strain

Viral stocks of USUV London were received from the Animal and Plant Health Agency (APHA) at a titre of 4 × 10^8^ PFU/ml. Virus stock was shipped on dry ice and was stored at -80 °C upon arrival. *Culex pipiens molestus* mosquito eggs were received from The Pirbright Institute [16] and were reared. To generate sufficient mosquitoes of a similar age for each experiment, mosquitoes were exposed to the Haemotek feeder overnight containing defibrinated horse blood (E & O Laboratories); and the following F1 generation were used for infection studies. The mosquito colony was maintained in the Insectary Facilities at Leahurst Campus, University of Liverpool.

### Oral infection of *Cx. pipiens molestus* with USUV

Mosquitoes were orally infected with a USUV titre of 4 × 10^7^ PFU/ml. A Haemotek feeder containing USUV and defibrinated horse blood was placed in the BugDorm cage for three hours. Mosquitoes were fed in the dark and non-bloodfed mosquitoes were removed. One hundred and thirty-eight *Cx. pipiens molestus* mosquitoes were infected with USUV at a titre of 4 × 10^7^ PFU/ml and incubated at 20 °C and 18 °C and 118 mosquitoes were infected and incubated at 22 °C. Following infection with USUV, mosquitoes were placed into DispoSafe pots and labelled with the number of days post infection on which they would be sampled.

### Time-series and incubation temperatures

*Culex pipiens molestus* mosquitoes were incubated at 18 °C, 20 °C and 22 °C using Sanyo TM MIR-154 incubators. Incubators were assessed prior to each experiment to check that the appropriate temperature could be maintained by the incubator. Incubation temperatures were monitored throughout experiments using a thermometer permanently placed in the incubator. Samples were collected immediately after feeding (0 days post-infection [dpi]) and 5, 10, 14, 17, 21 and 28 dpi.

### Forced salivation and sample collection

FlyNap (Blades Biological Limited, Kent) was used to anaesthetise the mosquitoes for forced saliva extraction. Once anaesthetised, mosquito proboscises were placed into a capillary tube containing mineral oil for 30 minutes. After 30 minutes, the mosquito bodies were placed into an Eppendorf tube containing 250 μl of Trizol (Thermo Fisher Scientific). The mosquito saliva was removed from the capillary tube using a micropipette and placed into a separate Eppendorf tube containing 100 μl of Trizol (Thermo Fisher Scientific).

### RNA extraction

Following placement of bodies and saliva into Trizol, RNA was extracted as per the manufacturer’s instructions. The RNA pellets was resuspended in 20 μl of nuclease-free water for 22 °C and 20 °C and 10 μl for 18 °C.

### Real-time reverse transcription PCR (RT-PCR)

Real-time RT-PCR was undertaken using a Roche 480 LightCycler (Roche, Basel, Switzerland). The total volume per well was 10 μl. This was made up of 5 μl of master mix, 0.4 μl of 10 μM forward (CGTGAAGGTTACAAAGTCCAGA) and reverse primers (TCTTATGGAGGGTCCTCTCTTC) targeting the nonstructural protein 1 gene, 0.2 μl of 10 μM probe, 1.9 μl of nuclease-free water, 0.5 μl of RT-mix and 2 μl of RNA template. RNA extract from a male *Cx. pipiens molestus* was used as a no-template control (NTC) and nuclease-free water was used as a negative control (NC). A reverse transcription step was undertaken at 50 °C for 30 minutes, followed by an initial activation step at 95 °C for 15 minutes. This was followed by 45 cycles of denaturation and combined annealing/extension at 94 °C for 15 seconds and 60 °C for 1 minute. USUV nucleic acid, SAAR 1776 strain (BEI resources, Bethesda, USA), was used as a positive control (PC). Saliva samples were run in triplicate, and whole body samples in duplicate on a 96-well real-time PCR plate. Half of the elution volume was used for 18 °C, in comparison to 20 °C and 22 °C to increase the sensitivity of this assay.

### Standard curve

Standards were produced by Integrated DNA Technologies using the NS1 sequence obtained from GenBank. Standards were serially diluted 1 in 10 from 1.7 × 10^12^ copies/μl to 1.7 × 10^2^ copies/μl, with a reaction efficiency of 1.832. Viral copy number was calculated using the standard curve for all samples with a crossing point value (Cp).

### Data analysis

Data was analysed using Rstudio [17]. Figures were produced using the ggplot and tidyverse packages [18, 19]. Survival analysis was undertaken using the ggsurvfit and survival packages [19, 20].

## Results

### Mortality rates and survival analysis for 22 °C, 20 °C and 18 °C

Three trends were observable in mortality rates identified in this experiment (Figure 1A); firstly, there was an increase in mortality over time for all temperatures conducted; secondly, an early peak in mortality followed by decline and then further increase was observed at all temperatures; and, thirdly, higher levels of mortality were observed at higher temperatures.

**Figure 1:**
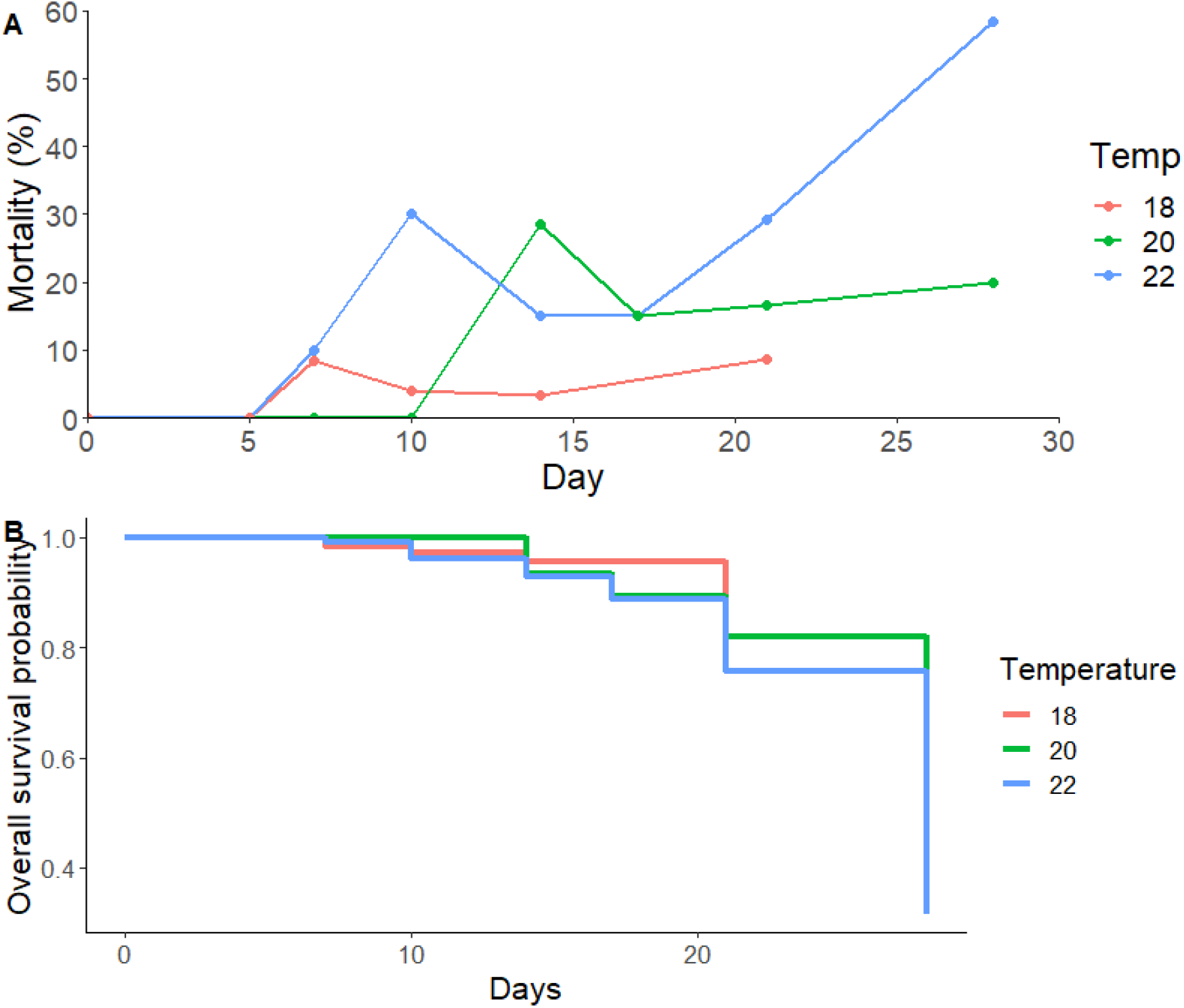
Total percentage mortality of *Culex pipiens molestus* at day of sampling (0, 5, 7, 10, 14, 21 and 28 days post infection). Mosquitoes were incubated up to 28 days for 22°C (n = 118) and 20 °C (n = 138), and up to 21 days for 18 °C (n = 138). Mosquito samples were collected at 0 dpi (n = 5 [22 °C]; n = 9 [20 °C]; n = 14 [18 °C]), 5 dpi (n = 5 [22 °C]; n = 10 [20 °C]; n = 10 [18 °C]), 7 dpi (n = 10 [22 °C]; n = 15 [20 °C]; n = 24 [18 °C]), 10 dpi (n = 10 [22 °C]; n = 14 [20 °C]; n = 25 [18 °C]), 14 dpi (n = 20 [22 °C]; n = 21 [20 °C]; n = 30 [18 °C]), 17 dpi (n = 20 [22 °C]; n = 20 [20 °C]), 21 dpi (n = 24 [22 °C]; n = 24 [20 ° C]; n = 35 [18 °C]) and 28 dpi (n = 24 [22 °C]; n = 25 [20 °C]). Percentage mortality was reported per DispoSafe pot in which infected mosquitoes were incubated (A). Kaplan-Meier plot demonstrating overall survival probability throughout experimental temperatures used. No left or right censoring occurred as all mosquitoes were used in analysis. Mosquitoes were incubated for up to 28 days for 22 °C and 20 °C, and for 21 days for 18 °C (B).

Using the same mortality data, the probability of survival at different time points was analysed with a Kaplan-Meier plot (Figure 1B). Survival analysis using the Cox Proportional Hazards model indicated that temperature had an effect on overall chance of survival, with statistical significance identified between 20 °C and 22 °C (p value <0.05).

### Proportion body and saliva positive

USUV RNA was detected in both bodies and saliva at 22 °C and 20 °C (Table 1). At both temperatures, 100% of mosquito bodies were positive at 0 dpi. Thereafter, the proportion positive dropped substantially, before starting to rise again. The rise started earlier and reached a higher level at 22 °C, compared to 20 °C. The time taken to reach the greatest proportion of body positive increased as the experimental temperature decreased, with 7 dpi, and 10 dpi being the time-point for greatest proportion of infected bodies at 22 °C and 20 °C, respectively.

**Table 1:** Number of positive and proportion positive mosquito saliva and body samples following oral inoculation with USUV and incubation at 22 °C and 20 °C for up to 28 days. The proportion of positive samples have been rounded to two decimal places. Data in graphical form can be found in Additional file 1: Figure A1.

| Temp (°C) | Day | Sample type | Positive samples | Total sampled | Proportion positive |
| --- | --- | --- | --- | --- | --- |
| 22 | 0 | Saliva | 0 | 5 | 0.00 |
| 22 | 0 | Body | 5 | 5 | 1.00 |
| 22 | 5 | Saliva | 0 | 5 | 0.00 |
| 22 | 5 | Body | 1 | 5 | 0.20 |
| 22 | 7 | Saliva | 4 | 9 | 0.44 |
| 22 | 7 | Body | 6 | 9 | 0.67 |
| 22 | 10 | Saliva | 1 | 7 | 0.14 |
| 22 | 10 | Body | 4 | 7 | 0.57 |
| 22 | 14 | Saliva | 1 | 17 | 0.06 |
| 22 | 14 | Body | 6 | 17 | 0.35 |
| 22 | 17 | Saliva | 0 | 16 | 0.00 |
| 22 | 17 | Body | 6 | 17 | 0.35 |
| 22 | 21 | Saliva | 0 | 17 | 0.00 |
| 22 | 21 | Body | 8 | 17 | 0.47 |
| 22 | 28 | Saliva | 1 | 10 | 0.10 |
| 22 | 28 | Body | 5 | 10 | 0.50 |
| 20 | 0 | Saliva | 0 | 9 | 0.00 |
| 20 | 0 | Body | 9 | 9 | 1.00 |
| 20 | 5 | Saliva | 1 | 10 | 0.10 |
| 20 | 5 | Body | 3 | 10 | 0.30 |
| 20 | 7 | Saliva | 0 | 15 | 0.00 |
| 20 | 7 | Body | 0 | 15 | 0.00 |
| 20 | 10 | Saliva | 0 | 14 | 0.00 |
| 20 | 10 | Body | 1 | 14 | 0.07 |
| 20 | 14 | Saliva | 0 | 15 | 0.00 |
| 20 | 14 | Body | 1 | 15 | 0.07 |
| 20 | 17 | Saliva | 0 | 17 | 0.00 |
| 20 | 17 | Body | 0 | 17 | 0.00 |
| 20 | 21 | Saliva | 0 | 20 | 0.00 |
| 20 | 21 | Body | 1 | 20 | 0.05 |
| 20 | 28 | Saliva | 0 | 19 | 0.00 |
| 20 | 28 | Body | 0 | 19 | 0.00 |

At both temperatures, 100% of mosquito saliva samples were positive at 0 dpi. At 22 °C, the proportion of saliva positives dropped to near zero by 5 dpi, then increased to a maxima at 7 dpi, with slow decline after this, before further increase at 28 dpi. At 20 °C, the percentage positive declined to zero by 7 dpi, with no further detection of USUV after 7 dpi.

### Relative viral copy number in mosquito bodies

The log of the viral copy number at each time point in mosquito bodies was compared to the log of the starting viral copy number (0 dpi) to determine how viral copy number changed throughout the experiment. At 20 °C, the log of the relative copy number decreased at each time point to be over three times smaller by 21 dpi. By contrast, the log of the relative copy number increased at 22 °C, albeit with evidence of a decrease between 7 dpi and 14 dpi. By 28 dpi, the log of the relative viral copy number was over three times higher than at the beginning of the experimental period (Figure 2). Relative viral copy number in saliva samples was not assessed as low numbers of positive saliva samples were observed with low viral copy numbers of USUV.

**Figure 2:**
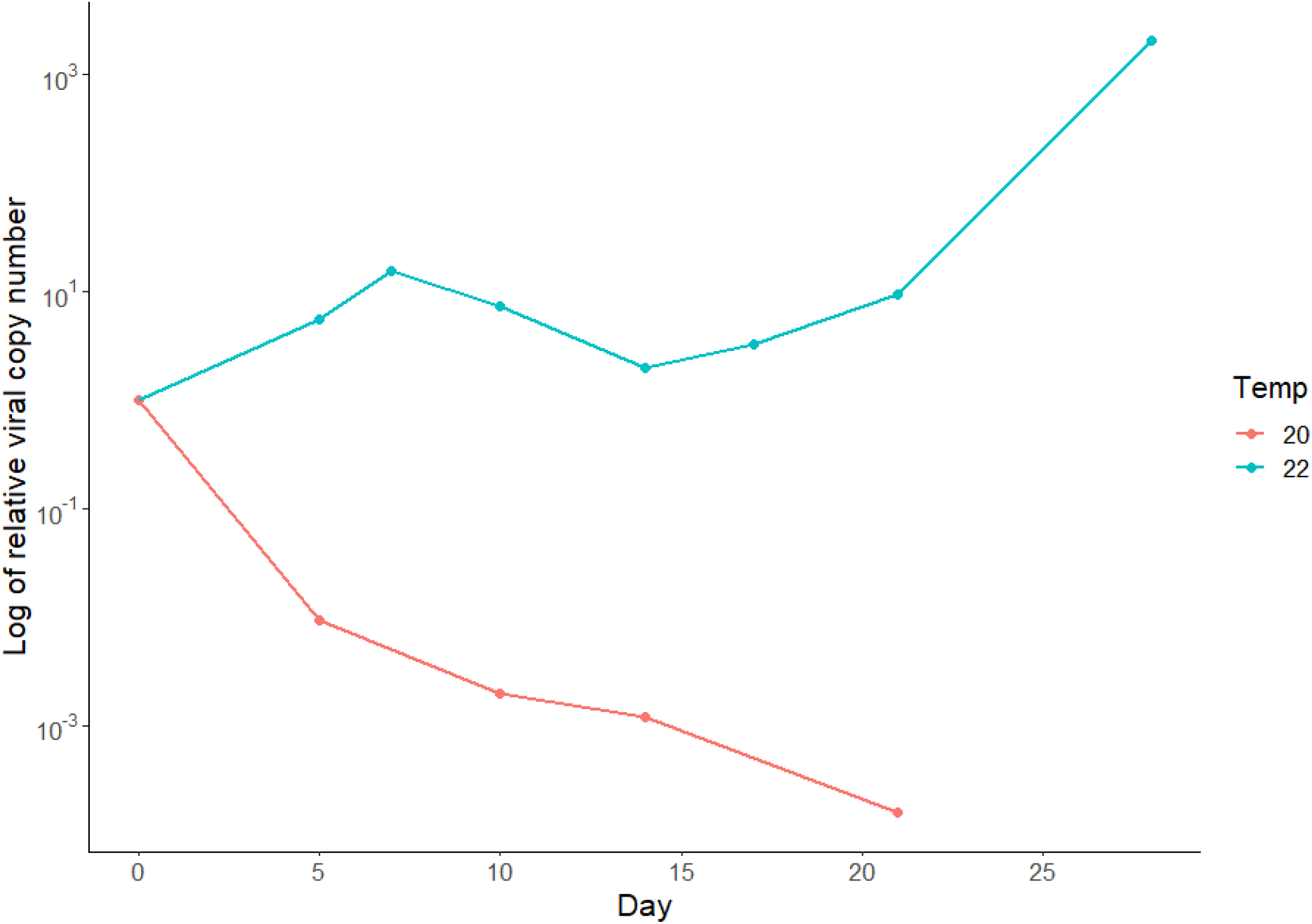
Change in viral copy number within mosquito bodies over time, relative to 0 dpi, at 22 °C and 20 °C. Data provided on log scale. Each datapoint represents the mean titre of positive mosquitoes at each timepoint. Pools of mosquitoes were fed spiked blood containing USUV at a titre of 4 × 10^7^ PFU/ml and were tested by real-time RT-PCR. Positive mosquito samples were assessed for viral titre at 0 dpi (n = 5/5^a^ [22 °C]; n = 9/9 [20 °C]), 5 dpi (n = 1/5 [22 °C]; n = 3/10 [20 °C]), 7 dpi (n = 6/9 [22 °C]; n = 0/15 [20 °C]), 10 dpi (n = 4/7 [22 °C]; n = 1/14 [20 °C]), 14 dpi (n = 6/17 [22 °C]; n = 1/15 [20 °C]), 17 dpi (n = 6/17 [22 °C]; n = 0/17 [20 °C]), 21 dpi (n = 8/17 [22 °C]; n = 1/20 [20 °C]) and 28 dpi (n = 5/10 [22 °C]; n = 0/19 [20 °C]). ^a^The numerator represents positive mosquito sample out of total number of mosquitoes tested at each timepoint.

### Study 2 (Temperature: 18 °C)

#### Proportion body and saliva positive

At 18 °C, and using a more sensitive detection method, all bodies were positive at 0 dpi, this then declined to a minimum at 7 dpi, before increasing again to 21 dpi (Table 2). All saliva samples were positive at 0 dpi, but the percent positive was zero at 5 dpi. As expected, the proportion of saliva positive was consistently less than the proportion of body positive. In comparison to 18 °C, USUV RNA was only detectable in saliva samples at 5 dpi in mosquitoes held at 20 °C but was detectable at 10, 14 and 21 dpi at 18 °C using the more sensitive assay.

**Table 2:** Number of positive and proportion positive mosquito saliva and body samples following oral inoculation with USUV and incubation at 18 °C for up to 21 days. The proportion of positive samples have been rounded to two decimal places. Data in graphical form can be found in Additional file 2: Figure A2.

| Day | Sample type | Positive samples | Total sampled | Proportion positive |
| --- | --- | --- | --- | --- |
| 0 | Saliva | 0 | 15 | 0.00 |
| 0 | Body | 15 | 15 | 1.00 |
| 5 | Saliva | 0 | 19 | 0.00 |
| 5 | Body | 6 | 19 | 0.32 |
| 7 | Saliva | 1 | 14 | 0.05 |
| 7 | Body | 1 | 13 | 0.07 |
| 10 | Saliva | 1 | 24 | 0.00 |
| 10 | Body | 2 | 24 | 0.08 |
| 14 | Saliva | 1 | 26 | 0.04 |
| 14 | Body | 4 | 29 | 0.14 |
| 21 | Saliva | 1 | 30 | 0.03 |
| 21 | Body | 6 | 30 | 0.21 |

#### Relative viral copy number in mosquito bodies

The relative viral copy number at 18 °C initially decreased between 0 and 5 dpi, then increased back to starting relative viral copy number at 14 dpi. A small decrease in relative viral copy number was then observed at 21 dpi (Figure 3). The experiment undertaken at 20 °C, as previously described, demonstrated a continuous decline in relative viral copy number throughout all time-points, with the lowest relative viral copy number occurring at 21 dpi. The fluctuation of relative viral copy number at 18 °C is more similar to 22 °C, although relative viral copy number at 22 °C increased to much higher levels than at the start of the experimental period.

**Figure 3:**
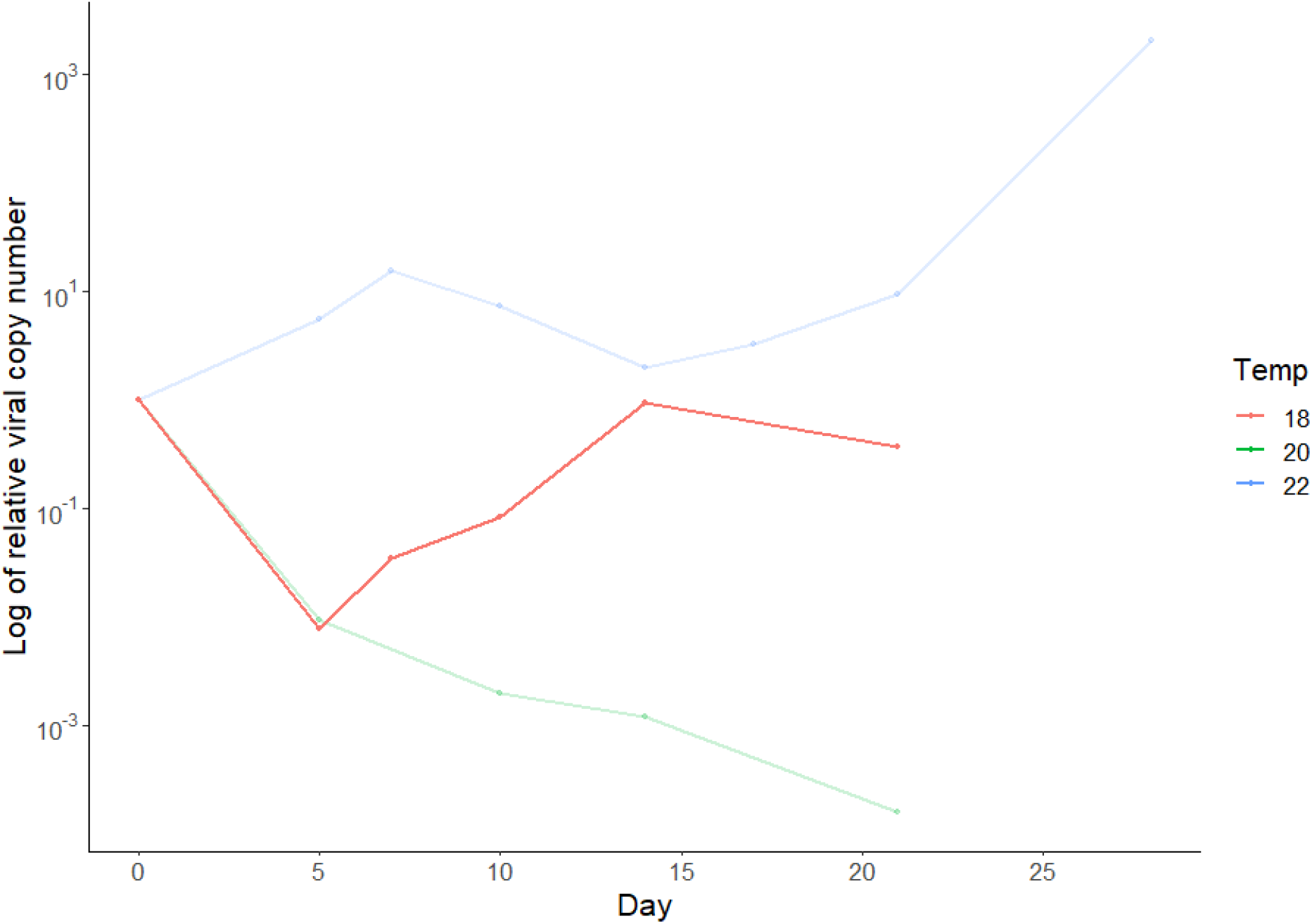
Demonstration of the log of the relative viral copy number within mosquito bodies over time, relative to day 0 viral copy number, at 20 °C and 18 °C. Data provided on log scale. Pools of mosquitoes were fed spiked blood containing USUV at a titre of 4 × 10^7^ PFU/ml and were tested by real-time RT-PCR. Each datapoint represents the mean titre of positive mosquitoes at each timepoint. Volume of elution buffer was halved for samples incubated 18 °C. Positive mosquito samples were assessed for viral titre at 0 dpi (n = 0/15^a^), 5 dpi (n = 6/19), 7 dpi (n = 1/13), 10 dpi (n = 2/24), 14 dpi (n = 4/29), 21 dpi (n = 6/30). ^a^The numerator represents positive mosquito sample out of total number of mosquitoes tested at each time point.

## Discussion

This study demonstrates the ability of a laboratory colony of *Cx. pipiens molestus* to become infected with, disseminate and be able to transmit the London strain of USUV under UK ambient temperatures. This has been achieved through detection of USUV RNA in mosquito bodies and saliva, with the greatest proportion of saliva positive samples occurring at 22 °C. This study suggests that temperatures in the range of 20 °C to 22 °C provide an important threshold for USUV replication. Additionally, we have demonstrated that temperatures of 19 °C and 20 °C, may facilitate transmission of the London strain of USUV [21]. The picture is more complex, however, as when a more sensitive detection method was used, the proportion of mosquitoes infected, and the number of viral genome copies, both increased even at the low temperature of 18 °C. Lower viral copy numbers can be potentially transmitted in mosquito saliva at cooler temperatures, and incorporation of a more sensitive assay in these experiments may lead to a reduction in false negative results. The strong effect of temperature on USUV infection supports that the vector competency and capacity of *Culex pipiens molestus* mosquitoes to transmit USUV may increase as temperatures rise. However, it is likely this relationship is not linear and that there may be a rise in extreme temperature which does not facilitate viral replication and may have detrimental effects on other elements required for successful pathogen transmission [22].

The data presented in this study produces contrasting results in comparison to another study which infected a laboratory colony of *Cx. pipiens* s.l. with the SAAR-1776 African strain of USUV. This study suggested a significant infection barrier of UK *Cx. pipiens* s.l. to this African strain. The difference in infection observed in these two studies may be suggestive of an evolutionary adaptation of USUV, London strain to be transmitted by *Cx. pipiens* s.l. mosquitoes in temperate regions. The evidence presented here suggests that successful viral replication can occur in mosquito bodies at 22 °C. USUV was detected in mosquito saliva at temperatures as low as 18 °C but viral titre in mosquito bodies at 20 °C and 18 °C did not exceed the viral titre at day 0, suggesting proliferation of USUV could not occur at temperatures less than 20 °C. Given higher relative viral copy numbers were identified at 18 ° C in comparison to 20 °C when a more sensitive assay was used, this highlights the importance of using sensitive assays for future studies focusing on USUV transmission in temperate regions.

This study has identified evidence of an eclipse phase in mosquito bodies – the subsequence decline of virus, as virus in the blood meal is digested or enters midgut cuts but is not yet replicating in the haemocoel or salivary glands. This is evidenced with a decrease in proportion positive and decrease in viral titre, followed by increase in both values at 22 °C and at 18 °C. At all temperatures, 100% of mosquito body samples were positive at 0 dpi; this is likely from residual virus in the mouthparts following a virus-spiked blood feed. Relative viral copy number in saliva samples was not assessed as low numbers of positive saliva samples were generated suggesting very low viral copy numbers of USUV in saliva samples. Relatively small sample sizes (five to 30 mosquitoes per time point) were sampled at each time-point in the various temperatures assessed. However, the number of infected mosquitoes sampled was nevertheless higher than in other infection studies with USUV [23].

The experimental model used here did not allow for fluctuations in temperature, as would occur naturally during the day and night. These temperature changes may have large consequences for viral replication and should be considered in future experiments. Effect of fluctuating temperatures on vector competency has been investigated with regards to West Nile virus (WNV) and *Cx. quinquefasciatus* and *Culex tarsalis* mosquitoes [24]. McGregor *et al* [24] showed that daily fluctuating temperature had a significant effect on viral titers in these mosquito species. Additionally, daily fluctuating temperatures has been shown to effect development time, fecundity and adult lifespan in *Culex pipiens molestus* [25]. Further work may also assess how behavioural effects of infected mosquitoes can impact on viral replication given that mosquitoes will identify desirable microclimates.

A real-time RT-PCR assay was used to detect USUV in mosquitoes but, as this assay detects fragments of genome, the viability of virus within samples tested could not be determined. The presence of viable virus would normally be demonstrated by plaque assay (with live virus killing cells, if present) but that was not undertaken here. Hence, despite finding positive USUV saliva by RT-PCR, it is not proven that there was the possibility of transmission of USUV at any temperature. Plaque assays were not undertaken because collected samples were placed immediately into Trizol, which kills live viruses. Nevertheless, there is a strong positive correlation of viral quantification through real-time RT-PCR and plaque assays for yellow fever virus (YFV), a related flavivirus to USUV, demonstrating some inference of infectivity may be possible when using real-time RT-PCR [26].

## Conclusion

This study has demonstrated the vector competency of a laboratory colony of *Cx. pipiens molestus* for the London strain of USUV, an African lineage 3 strain. Here, we demonstrate a critical temperature threshold between 22 °C and 20 °C, demonstrating active USUV replication and potential transmission at 22 °C. We highlight the possibility of permissiveness of transmission of USUV in London and, more broadly, the south-east of England. We highlight the importance of assay sensitivity when performing vector competency studies at cooler temperatures. Furthermore, preliminary data has been provided in this study which can be used to inform future infection studies which have the aim of extracting extrinsic incubation period (EIP) information to inform epidemiological and disease-risk models. We suggest that sustained transmission is possible at current ambient temperatures, and with the consideration of climate change, it is possible the areas of risk of USUV transmission in the UK may expand in coming years.

## Supplementary information

Additional file 1: Figure A1: Proportion of body and saliva positive samples at 22 °C and 20 °C. Saliva samples at 0 dpi were positive, likely from virus contamination in the mouthparts, and so here are forced to one (A). Body samples at 0 dpi were positive, likely from virus in the bloodmeal (B). Pools of mosquitoes were fed spiked blood containing USUV at a titre of 4 × 10^7^ PFU/ml and were tested by real-time RT-PCR.

Additional file 2: Figure A2: Proportion of body positive and saliva positive samples at, 22 °C, 20 °C and 18 °C. Samples incubated at 18 °C were analysed using an elution buffer at half the volume of samples at 20 °C. Pools of mosquitoes were fed spiked blood containing USUV at a titre of 4 × 10^7^ PFU/ml and were tested by real-time RT-PCR. Saliva samples at 0 dpi were taken immediately after blood feeding and residual virus present in mouthparts likely contaminated saliva as it was expectorated giving rise to false positive results (A). Here, the proportion of saliva positive samples is forced to one. Body samples at 0 dpi contained virus in the blood meal and so the proportion of body positive samples was 1 at 0 dpi (B).

## Abbreviations

Dpi: days post infection
EPI: extrinsic incubation period
RT-PCR: real-time reverse transcription polymerase chain reaction
USUV: Usutu virus
WNV: West Nile virus

## Declarations

## Acknowledgements

We would like to thank The Pirbright Institute for provision of the laboratory colony of *Cx. pipiens molestus*.

## Author’s contributions

NS was responsible for the primary data collection, analysis and manuscript production. JP assisted with primer design, data collection and analysis. KS provided technical support in colony rearing and sample collection. JTH assisted with RNA extractions. MBl, GH and MBa assisted with development of the experimental protocol. NJ provided provisions of viral stocks. All authors have read and approved the final manuscript.

## Funding

This research was funded by the National Institute for Health & Social Care Research Health Protection Research Unit (NIHR) in Emerging and Zoonotic Infections at the University of

Liverpool in partnership with the UK Health Security Agency (UKHSA), in collaboration with Liverpool School of Tropical Medicine and The University of Oxford.

## Availability of data materials

The datasets generated and/or analysed during the current study are available from the corresponding author upon reasonable request.

## Ethical approval and consent to participate

Not applicable

## Consent for publication

Not applicable

## Author details

Department of Livestock and One Health, Institute of Infection, Veterinary and Ecological Sciences, University of Liverpool, UK

