## Additional file 2 for "Impact of temperature on vector competence of *Culex pipiens molestus*: implications for Usutu virus transmission in temperate regions"

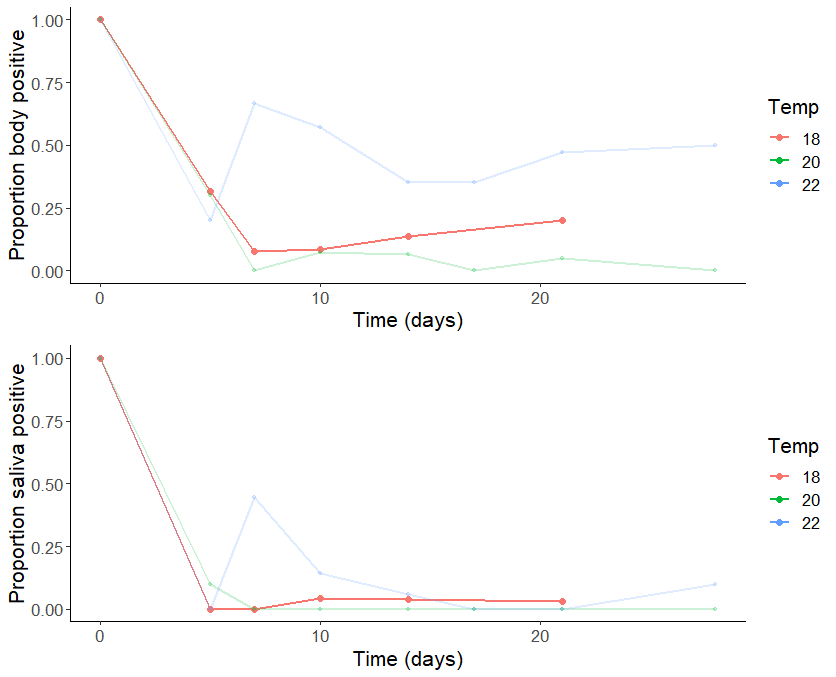

Figure A2: Proportion of body positive and saliva positive samples at, 22 ˚ C, 20 ˚ C and 18 ˚ C. Samples incubated at 18 ˚ C were analysed using an elution buffer at half the volume of samples at 20 ˚ C. Pools of mosquitoes were fed spiked blood containing USUV at a titre of 4 x 10^7^ PFU/ml and were tested by real-time RT-PCR. Saliva samples at 0 dpi were taken immediately after blood feeding and residual virus present in mouthparts likely contaminated saliva as it was expectorated giving rise to false positive results (A). Here, the proportion of saliva positive samples is forced to one. Body samples at 0 dpi contained virus in the blood meal and so the proportion of body positive samples was 1 at 0 dpi (B). Mosquito samples were tested at 18 ˚ C are as follows: 0 dpi (n = 15); 5 dpi (n = 19); 7 dpi (n = 13); 10 dpi (n = 24); 14 dpi (n = 29); and 21 dpi (n = 30).
