## Additional file 1 for "Impact of temperature on vector competence of *Culex pipiens molestus*: implications for Usutu virus transmission in temperate regions"

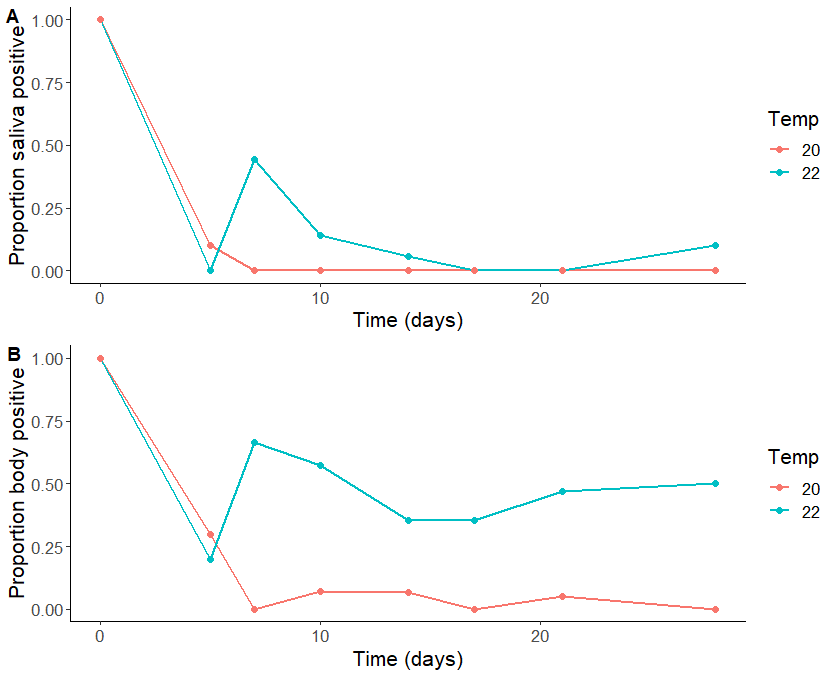

Figure A1: Proportion of body and saliva positive samples at 22 ˚ C and 20 ˚ C. Saliva samples at 0 dpi were positive, likely from virus contamination in the mouthparts, and so here are forced to one (A). Body samples at 0 dpi were positive, likely from virus in the bloodmeal (B). Pools of mosquitoes were fed spiked blood containing USUV at a titre of 4 x 10^7^ PFU/ml and were tested by real-time RT-PCR. Mosquito samples were collected at 0 dpi (n = 5 [22 ˚ C]; n = 9 [20 ˚ C]), 5 dpi (n = 5 [22 ˚ C]; n = 10 [20 ˚ C]) , 7 dpi (n = 9 [22 ˚ C]; n = 15 [20 ˚ C]), 10 dpi (n = 7 [22 ˚ C]; n = 14 [ 20 ˚ C]), 14 dpi (n = 17 [22 ˚ C]; n = 15 [20 ˚ C]), 17 dpi (n = 17 [22 ˚ C]; n = 17 [20 ˚ C]), 21 dpi (n = 17 [22 ˚ C]; n = 20 [20 ˚ C]) and 28 dpi (n = 10 [22 ˚ C]; n = 19 [20 ˚ C]).
